# Medin-Induced Pro-inflammatory and Prothrombotic Activation of Coronary Artery Endothelial Cells: A Potential Novel Mediator Linking Aging and Atherosclerosis

**DOI:** 10.64898/2026.07.02.736227

**Authors:** Kaleb Morrow, Nina Karamanova, Randy Woltjer, Victoria Krajbich, Jingmin Shu, Ming Li, Chengyun Tang, Alana Maerivoet, Jillian Madine, Yabing Chen, Raymond Q. Migrino

## Abstract

**Background:** Age is the most important risk factor for coronary artery disease (CAD) independent of traditional risk factors. Aging induces classic pro-inflammatory and prothrombotic vascular phenotypic changes whose molecular mediators remain poorly understood. Medin is a common cleavage product protein that accumulates in vasculature with aging and shown to cause endothelial dysfunction. Its role in CAD is unknown. The study aimed to evaluate the effects of medin on human coronary artery endothelial cell (HCAEC) pro-inflammatory and prothrombotic activation and establish the relationship between medin and coronary atherosclerosis in human decedents.

**Methods:** HCAECs were exposed to physiologic dose of medin (5 µM) for 20 hours and ribonucleic acid sequencing (RNAseq) with signaling pathway analyses and reverse transcription polymerase chain reaction of select pro-inflammatory and prothrombotic genes performed. Corresponding protein expression was measured by Western blot or enzyme linked immunosorbent assay in HCAECs exposed to medin (5 µM) without or with nuclear factor-κB (NFκB) inhibitor RO106-9920 (10 µM). Coronary arteries from 40 deceased individuals underwent immunohistochemistry and medin and plaque burden were quantified and their relationship evaluated.

**Results:** RNAseq showed predominant pro-inflammatory gene expression changes induced by medin. HCAECs treated with medin showed increased phosphorylated NFκB, elevated protein expression of interleukin (IL)-6, IL-8, monocyte chemotactic protein (MCP)-1, intercellular adhesion molecule (ICAM)-1, vascular cell adhesion molecule (VCAM)-1 and plasminogen activator inhibitor (PAI)-1 and reduced protein expression of thrombomodulin; these changes were reversed by RO106-9920 co-treatment. In human tissues, coronary artery medin strongly correlated with plaque burden (R=0.76, p<0.0001) and coronary macrophage content (R=0.72, p<0.0001). Coronary arteries from decedents with myocardial infarction had higher medin than those without (5.53±2.67% versus 0.02±0.02%, p=0.0005).

**Conclusions:** Medin induced NFκB-mediated endothelial cell pro-inflammatory and prothrombotic activation and was strongly associated with coronary plaque burden and inflammation. Medin is a novel candidate mediator linking aging and coronary atherosclerosis.

## INTRODUCTION

Coronary artery disease (CAD) is the leading cause of death worldwide^1^ and age is the most important risk factor for CAD^2^. Aging, even in the absence of traditional cardiovascular risk factors (such as smoking, hyperlipidemia, diabetes and hypertension), leads to classic vascular aging phenotype that includes arterial inflammation, endothelial and smooth muscle dysfunction, prothrombotic milieu, calcification and stiffening^3^. Aging is associated with chronic low grade inflammation which predisposes the vasculature to the development of atherosclerosis^3^. Chronic upregulation of pro-inflammatory mediators such as interleukin (IL)-6, IL-1β and tumor necrosis factor (TNF)-α are induced during the aging process through activation of many pro-inflammatory signaling pathways including the nuclear factor-κB (NFκB) signaling pathway^4^.Advancing age is also associated with increased rates of arterial and venous thrombosis^5^. The frequency of myocardial infarction and stroke increases significantly with aging with about 80% of all fatal myocardial infarctions occurring in the elderly^6^. Aging is associated with an alteration in the fine balance between pro- and anti-thrombotic components. Hypercoagulability in the blood of the elderly results from elevation of plasma coagulation factors VII, VIII and XIII, plasminogen activator inhibitor (PAI)-1 without accompanying rise in natural anticoagulant factors such as proteins C and S, shifting the balance to a pro-thrombotic milieu^7^. The endothelium normally provides an antithrombotic surface through expression of thrombomodulin and tissue plasminogen activator^8^. Thrombomodulin is a 100 kDa transmembrane glycoprotein expressed in abundance by vascular endothelial cells and is a critical component of the anticoagulant protein C pathway^9^.

Although the molecular, functional and structural changes of vascular aging and atherosclerosis have been documented^2,3,10^, the agents mediating these changes remain largely unknown. Recently, the most common human amyloidogenic protein^11^, medin, which accumulates in the vasculature with aging^12^ was shown to accumulate and cause endothelial dysfunction in human cerebral arteries^13–15^ while inducing pro-inflammatory activation of endothelial cells through NFκB-dependent signaling pathway^15–17^. Medin is a 50 amino acid fragment from parent protein milk factor globule-E8 (MFGE8)^11^. Vascular accumulation in cerebral arteries is associated with Alzheimer’ disease, vascular dementia and aging-associated cerebrovascular disease^13,18–20^. Originally discovered as the major component of aortic amyloid, medin is associated with aortic aneurysm and dissection^21,22^. While medin has been known to accumulate in coronary arteries^12^ its role in CAD remains unknown. We aim to test the effects of medin on human coronary artery endothelial cells (HCAECs) focusing on pro-inflammatory and prothrombotic mediators, establish the relationship between medin and coronary atherosclerosis in deceased human subjects and identify potential mechanistic basis for the relationship.

## METHODS

### Recombinant Medin

pOPINS-medin was expressed in Lemo 21 (DE3) and purified using a Ni^2+^-NTA column with SUMO protease I cleavage as described in prior work^15^. Purity of medin was confirmed to be >95% by SDS-PAGE, snap frozen and stored at −80°C. *Limulus* Amebocyte Lysate assay (Pierce, Dallas TX) was used to confirm that endotoxin levels were <0.5 ng/mL.

### Treatment Conditions for HCAECs

HCAECs (Cell Systems; Kirkland, WA, passages 6-10) were seeded to 90% confluence. For ribonuclear acid sequencing (RNAseq) assay, HCAECs were treated with vehicle or medin 5 µM (dose selected from prior observation that human aortic tissue medin content range was 0-13.72 µM)^15^ for 20 hours followed by cell harvest. For gene expression assays, HCAECs were treated with 0, 0.5, 1 and 5 µM of medin for 20 hours and cell lysates were collected. For protein expression assays, HCAECs were treated with vehicle, medin 5 µM without or with NFκB inhibitor RO106-9920 (Tocris Biosciences, Bristol United Kingdom, 10 µM) or RO106-9920 10 µM alone for 20 hours. RO106-9920 dose was based on prior study demonstrating inhibitor effectiveness in suppressing inflammatory activation of human umbilical vein endothelial cells treated with medin^15^. To confirm that the biological effects of medin are due to its structural composition and not a random protein effect, in separate experiments, HCAECs were treated with vehicle, medin 5 µM and scrambled medin 5 µM (custom made by Genscript, Piscataway New Jersey, Sequence: SVYLQDQWNNQVQLSRDEVALGIKAKRFGQTNGWSGAGFNTSIDGVAGFV) and gene expression outcomes were compared.

### RNA Sequencing and Bioinformatics Analyses

Total RNA was isolated from four biological replicates of each experimental group (vehicle and medin 5 µM). Strand-specific mRNA libraries were constructed using a dUTP-based protocol. Briefly, poly(A)+ mRNA was enriched using oligo(dT)-conjugated magnetic beads, fragmented, and reverse-transcribed to generate first-strand cDNA. Second-strand cDNA synthesis was performed with dUTP incorporation to preserve strand information. Following end repair, adapter ligation, and PCR amplification, libraries were converted into DNA nanoballs (DNBs). Libraries were sequenced on the DNBSEQ platform (BGI, Shenzhen, China) using a paired-end 150-bp (PE150) strategy. An average of 22.07 million clean reads and 6.62 Gb of clean bases were generated per sample. Sequencing quality was high, with mean Q20 and Q30 scores exceeding 97% and 91%, respectively, and a mean clean-read ratio greater than 97%.

Raw sequencing reads were processed using SOAPnuke (v2.3) to remove adapter-contaminated^23^, low-quality, and ambiguous reads. Clean reads were aligned to the human reference genome (GRCh38.p14) using HISAT2 (v2.0.4)^24^. Alignment quality was evaluated based on mapping statistics, transcript coverage, and sequencing saturation. Gene and transcript abundances were quantified using Bowtie2 (v2.2.5)^25^ and RSEM (v1.2.8)^26^. After filtering and alignment, an average of 98.85% of clean reads mapped to the reference genome, with 86.97% aligning to annotated gene regions. In total, 17,355 genes were detected across all samples, demonstrating high sequencing quality and sufficient transcriptome coverage for downstream analyses.

Differential gene expression analysis was performed using the DESeq2 package in R, which models count data using a negative binomial distribution^27^. Genes with an FDR-adjusted *P* value ≤ 0.05 and a log2 fold change (log2FC) < −1 or > 0.58, corresponding to at least a 50% decrease or increase in expression, respectively, were considered differentially expressed.

Functional enrichment analyses of differentially expressed genes (DEGs) were conducted using Gene Ontology (GO) and Kyoto Encyclopedia of Genes and Genomes (KEGG) databases^28^. Candidate genes were mapped to GO terms and KEGG pathways, and enrichment significance was assessed using a hypergeometric test implemented through the R function *phyper*. Multiple-testing correction was performed using the Bioconductor *qvalue* package to estimate false discovery rates^29^. GO terms and KEGG pathways with Q values ≤ 0.05 were considered significantly enriched.

KEGG pathway enrichment results were visualized using bubble plots generated with ggplot2. The Rich Ratio, defined as the ratio of enriched DEGs to the total number of genes annotated within a given pathway, was plotted on the x-axis, while enriched pathways were displayed on the y-axis. Bubble size represented the number of enriched genes, and bubble color corresponded to the enrichment Q value. For pathway-level visualization, gene symbols were mapped to Entrez Gene identifiers using the org.Hs.eg.db annotation package^30^. Genes lacking valid Entrez annotations were excluded, and duplicate Entrez IDs were collapsed by averaging log2 fold-change values. The Pathview package (Bioconductor) was used to integrate gene expression changes into KEGG pathway diagrams^31^. For visualization of the TNF signaling pathway (KEGG pathway hsa04668)^32^, log2 fold-change values from the HCAEC_2 (medin-treated) versus HCAEC_1 (vehicle) comparison were overlaid onto the KEGG reference map. Upregulated and downregulated genes were displayed using the default Pathview color scale, with red and green indicating increased and decreased expression, respectively.

### Gene Expression Assays

Total RNA samples from the HCAEC cell lysates were prepared using the Aurum Total RNA kit (cat#732-6820) and iScript cDNA Synthesis Kit (cat# 170-8890) was used for cDNA preparation. All kits obtained from BIO-RAD Laboratories, Inc. CA. Gene expression was measured using RT PCR (PAI1 Primers obtained from Sigma-Aldrich Corp. The Woodlands, TX. Other primers obtained from IDT DNA Technologies, Coralville IL) with β-Actin (ACTB) as the reference normalization gene. Primer used were: ACTB PrimeTime qPCR Primers, Exon Location:1-2, RefSeq Number: NM_001101 (Primer1: 5’-ACA GAG CCT CGC CTT TG-3’ Primer 2: 5’-CCT TGC ACA TGC CGG AG-3’), IL-8 PrimeTime qPCR Primers, Exon Location: 3-4, RefSeq Number: NM_000584 (Primer1: 5’-TGT CTG GAC CCC AAG GAA-3’ Primer2: 5’-CAT CTT CAC TGA TTC TTG GAT ACC-3’), IL-6 F:5’- AAC CTG AAC CTT CCA AAG ATG-3’ R:5’-TCT GGC TTG TTC CTC ACT AC −3’, IL1-b F:5’- CAG CTA CGA ATC TCC GAC CAC −3’ R:5’-GGC AGG GAA CCA GCA TCT TC-3’, ICAM1 PrimeTime qPCR Primers, Exon Location: 2-3, RefSeq Number: NM_000201(Probe 5’-/56-FAM/AGTCCAGTA/ZEN/CACGGTGAGGAAGGT/3IABkFQ/3’ Primer1: 5’-GCTATTCAAACTGCCTGATG-3’ Primer2: 5’-GCGTAGGGTAAGGTTCTTGC-3’), VCAM1 PrimeTime qPCR Primers, Exon Location: 7-8, RefSeq Number: NM_001199834 (Primer 1: 5’-AAG CAT GTC ATA TTC ACA GAA CTG-3 Primer 2: 5’-AAC CCA AAC AAA GGC AGA GTA-3’) PAI-1 SENS: 5’-CTC TCT CTG CCC TCA CCA AC-3’ ANTIS: 5’-GTG GAG AGG CTC TTG GTC TG TG-3’, MCP-1 (CCL2) PrimeTime qPCR Primers, Exon Location: 1–2, RefSeqNumber: NM_002982 (Primer 1: 5’ – AGCAGCCACCTTCATTCC-3’ Primer 2: 5’-GCCTCTGCACTGAGATCTTC-3’) and TNF-α PrimeTime qPCR Primers, Exon Location:1b-4a, RefSeqNumber: NM_000594 (Primer 1: 5’-TGCACTTTGGAGTGATCGG-3’ Primer 2: 5’-TCAGCTTGAGGGTTTGCTAC-3’).

### Protein Expression Assays

HCAEC protein expression were obtained either by Western blot (WB) or enzyme linked immunosorbent assay (ELISA). WB was performed using antibodies to D141/Thrombomodulin rabbit mAb (E7Y9P; Cell Signaling Technologies, Danvers, MA) diluted 1:1000, PAI-1 Rabbit mAb (D9C4; Cell Signaling Technologies, Danvers, MA) diluted 1:1000, phospho-NFκB p65 (ser536) 93H1 (rabbit mAb, Cell Signaling#3033, 1:1000), total NFκB p65 D14E12XP (rabbit mAb Cell Signaling#8242, 1:1000). IRDye 680RD/800CW 2° Abs (Li-Cor; Lincoln, NE) were used in a 1:10000 dilution and imaged on the LI-COR Odyssey Imaging System (Li-Cor; Lincoln, NE). Results were normalized to β-actin control (8H10D10 mouse mAb, Cell Signaling#3700, 1:1000). ELISA for IL-6, IL-8 and MCP-1 in conditioned media was performed using DuoSet ELISAs (R&D Systems; Minneapolis, MN). ELISA for ICAM-1 and VCAM-1 in total cell lysate was performed using DuoSet ELISAs (R&D Biosystems; Minneapolis, MN).

### Activated Protein C Assay

HCAEC (Cell Systems; Kirkland, WA) were seeded in 96 well plates to 90% confluence. The cells were treated in duplicate for 20 hours at 37°C, 5% CO_2_ with either vehicle or recombinant Medin 5uM in Normal Glucose Media (Cell Systems; Kirkland, WA). After treatment, the cells were rinsed with D-PBS and incubated in Opti-MEM (Gibco; Brooklyn, NY) at 37°C, 5% CO_2_ for 1 hour. Activated Protein C was measured using a modification of the protocol from Hultstrom et al.^33^. 50µL of APC Buffer containing 0.1% BSA, 3mM CaCl_2_, 0.6mM MgCl_2_, 8.28µg/mL Human Protein C (Prolytix; Essex Junction, VT), and 61.26 ng/mL Human α-Thrombin (Prolytix; Essex Junction, VT) in Ca^2+^/Mg^2+^ free PBS was added to each well. Soluble Thrombomodulin (R&D Biosystems; Minneapolis, MN) was added to the APC Buffer for Medin 5µM treated cells in 3 different doses: 0.5µM, 1µM, and 1.5µM. Cells were incubated at 37°C, 5% CO_2_ for 30 minutes before adding 10uL of 2 U/mL Hirudin (Sigma-Aldrich; St. Louis, MO) to each well. Cells were incubated for an additional 10 minutes at 37°C, 5% CO_2_ to terminate the reaction. 40uL of 2.5mg/mL Activated Protein C Chromogenic Substrate (Anaira Diagnostica; West Chester, OH) was then added to each well. The plate was put into a SpectraMax 190 plate reader (Molecular Devices; San Jose, CA) and absorbance was read every 15 seconds for 2 hours at 405nm and 37°C with shaking between reads.

### Human Coronary Tissues

Coronary tissues were obtained from 14 donor participants of the Banner Sun Health Research Institute Brain and Body Donation Program, a longitudinal clinicopathologic study of aging and neurodegenerative disease around retirement communities in northwest metropolitan Phoenix Arizona region (Dr. Thomas Beach, Director)^34^. The program was approved by the Sun Health Research Institute and written informed consent was obtained from all subjects or their legal representatives. Deidentified coronary tissue sections were shared with the Phoenix VA under a Materials Transfer Agreement approved by the Phoenix VA Institutional Review Board and Research and Development Committees.

For external validation of our findings, a separate cohort of coronary tissues were studied. Deidentified coronary tissues from 26 deceased individuals at the Oregon Health and Science University underwent histologic examination. Consent for research use of tissues was obtained from the legal next of kin prior to autopsy. Use of deidentified postmortem human tissue is not considered human subjects research by the OHSU Institutional Review Committee and does not require specific approval.

### Coronary Artery Histology

Immunohistochemistry was performed at the Phoenix VA on 5 uM thick human heart tissue sections from paraffin blocks and antigen retrieval was performed using mouse monoclonal specific anti-Medin antibody (Prothena 18G, 1:1000, generously provided by Prothena Biosciences, San Francisco CA) followed by diaminobenzidine (DAB) staining (Vector Laboratories DAB Substrate Kit, Peroxidase with Nickel, SK-4100) using methods previously described^35^. After scanning slides on the MoticEasyScan Digital Slide Scanning System with a high-precision motion control and high-resolution digital imaging systems, slides were soaked in Xylene overnight with agitation to remove coverslips. The tissue was rehydrated by processing slides in Xylene 2 cycles, then in graded alcohols to distilled water (2x100%,1x95%,1x70% EthOH, then 2xddH2O) followed by staining with Movat Pentachrome Stain (Modified Russel-Movat, kit ab245884). A contiguous coronary section was then selected, and immunohistochemistry was performed using mouse monoclonal anti-human CD68 antibody (Clone KP1, macrophage marker, Dako Agilent, Santa Clara CA, 1:80 dilution).

Images were analyzed using QuPath Open Software for Bioimage Analysis version 0.6.0^36^. The two largest epicardial coronary arteries on the tissue section were chosen for analysis in 12 donors; in 2 donors, only one coronary artery was available for analysis in the tissue section. The outlines of the coronary lumen, the internal elastic lamina (separating the tunica intima and tunica media) and the external elastic lamina (separating the tunica media from the adventitia) were traced. The tunica intima is composed of both the intima and the neointima and the area of the tunica intima is designated as plaque area in this study. The intima-media is composed of both the tunica intima and tunica media and the intima-media area is the area enclosed by the luminal border and the external elastic lamina. Plaque area % was calculated as (plaque area/intima-media area)X100%. Coronary medin and CD68 were quantified using threshold method^36^. Briefly, a section of the coronary arteries with stained and unstained regions were selected and stain vectors were analyzed by the software. Pixel classification was done by setting a threshold pixel value that delineates stained versus unstained region. The software provided the sum of the areas of regions with stain. Medin burden in the intima-media region was quantified from the DAB stain and expressed as medin area %, calculated as (medin area/intima-media area)X100%. Macrophage burden in the intima-media region was quantified from the DAB stain and expressed as CD68% area, calculated as (CD68 area/intima-media area)X100%. Because of the variability of the extent of the coronary adventitia, this area was excluded in the measurement of coronary area, medin and CD68 burden.

A separate external validation cohort was done to confirm the relationship between coronary medin and plaque burden using tissues from 26 decedents at the Oregon Health and Science University (OHSU). Separate DAB immunostaining was done at OHSU using anti-medin antibody (Prothena 18G, 1:2000 dilution) developed using horseradish peroxidase (Vector ELITE HRP kit) and hematoxylin and eosin staining as per prior methods^35^. Quantification of medin area % and plaque area % were done using similar methods described above using QuPath software. For the OHSU cohort, only 1 epicardial coronary artery section was analyzed for each decedent.

### Data and Statistical Analyses

Data are expressed as mean±standard error of means with significant p-value set at p<0.05 (two-sided). HCAECs outcome readouts, coronary medin area, plaque area and CD68 area were analyzed using one-way analysis variance (ANOVA) with pairwise Tukey’s multiple comparison test. When there was heavy-tailedness in the outcome distribution, natural log transformation was done and transformed values were used for the analyses. If heavy-tailedness persisted with transformed values, Friedman test was performed on raw data and pairwise Dunn’s multiple comparison test was done. Student’s t-test was used to compare medin and CD68 content between MI versus no MI coronary arteries. Correlation was performed using Spearman analyses. Analyses were performed using GraphPad Prism 10 (GraphPad Software, Boston MA). RQM had full access to all data in the study and takes full responsibility for their integrity and data analyses.

## RESULTS

### HCAEC RNA sequencing

Table 1A shows the top 10 genes upregulated in HCAECs following medin treatment. The list is dominated by pro-inflammatory genes including VCAM-1, interferon gamma-induced protein 10 and granulocyte chemotactic protein 2. Figure 1A shows signaling pathway analyses confirming that pro-inflammatory pathways had the highest gene expression changes with TNF, advanced glycation endproduct-receptor for advanced glycation endproduct (AGE-RAGE) and NFκB signaling pathways among the most altered by medin treatment. Figure 1B shows specific genes differentially expressed in the TNF signaling pathway that informed our selection of genes for rtPCR confirmation.

**Table 1.** Top 10 differentially expressed genes in HCAECs following medin treatment.

| Gene ID | Symbol | Protein Encoded | log2 | Qvalue |
| --- | --- | --- | --- | --- |
| **A. Overexpressed** |  |  |  |  |
| 7412 | 'VCAM1' | Vascular cell adhesion molecule 1 | 9.73 | 7.78E-18 |
| 3627 | 'CXCL10' | Interferon gamma-induced protein 10 | 8.93 | 2.40E-13 |
| 6372 | 'CXCL6' | Granulocyte chemotactic protein 2 | 8.86 | 2.04E-108 |
| 4600 | 'MX2' | MX dynamin-like GTPase2 | 8.77 | 1.19E-07 |
| 129607 | 'CMPK2' | Cytidine/uridine monophosphate kinase 2 | 7.98 | 2.99E-10 |
| 969 | 'CD69' | Cluster of differentiation 69 | 7.95 | 6.22E-10 |
| 1437 | 'CSF2' | Granulocyte-macrophage colony-stimulating factor | 7.87 | 1.36E-33 |
| 10537 | 'UBD' | Ubiquitin D | 7.58 | 2.64E-10 |
| 684 | 'BST2' | Tetherin | 6.90 | 7.60E-14 |
| 2921 | 'CXCL3' | C-X-C motif chemokine 3 | 6.86 | 2.22E-42 |
| **B. Underexpressed** |  |  |  |  |
| 283106 | 'CSNK2A3' | Casein kinase 2 alpha 3 | -22.19 | 5.07E-09 |
| 613227 | 'HIGD1C' | Hypoxia-inducible domain family member 1C | -3.34 | 0.005 |
| 4741 | 'NEFM' | Neurofilament medium chain | -3.28 | 0.006 |
| 401190 | 'RGS7BP' | Regulator of G protein signaling 7 binding protein | -3.00 | 4.47E-06 |
| 1393 | 'CRHBP' | Corticotropin releasing hormone binding protein | -2.93 | 0.002 |
| 112268293 | 'TMA7B' | Translation machinery associated 7 homolog | -2.88 | 0.004 |
| 5042 | 'PABPC3' | Polyadenylate-binding protein 3 | -2.88 | 0.006 |
| 51206 | 'GP6' | Glycoprotein VI | -2.27 | 2.76E-11 |
| 2124 | 'EVI2B' | Ecotropic viral integration site 2B | -2.24 | 1.50E-04 |
| 9865 | 'TRIL' | TLR4 interactor with leucine rich repeats | -2.21 | 2.25E-07 |
| **C. Atherothrombosis**<br>**DEG Genes** |  |  |  |  |
| 7412 | 'VCAM1' | Vascular cell adhesion molecule 1 | 9.73 | 7.78E-18 |
| 6401 | 'SELE' | E-selectin | 6.81 | 4.86E-05 |
| 3553 | 'IL1B' | Interleukin-1β | 6.77 | 1.37E-07 |
| 3383 | 'ICAM1' | Intercellular adhesion molecule 1 | 4.54 | 1.87E-39 |
| 6347 | 'CCL2' | Monocyte chemoattractant protein 1 | 4.22 | 2.85E-134 |
| 4791 | 'NFKB2' | Nuclear factor kappa B subunit 2 | 2.46 | 1.80E-33 |
| 5054 | 'SERPINE1' | Plasminogen activator inhibitor-1 | 1.58 | 1.06E-25 |
| 7056 | 'THBD' | Thrombomodulin | -1.74 | 7.49E-22 |
| 4846 | 'NOS3' | Nitric oxide synthase 3 | -1.24 | 1.12E-04 |
| 5327 | 'PLAT' | Tissue-type plasminogen activator | -0.73 | 9.68E-16 |

**Figure 1.**
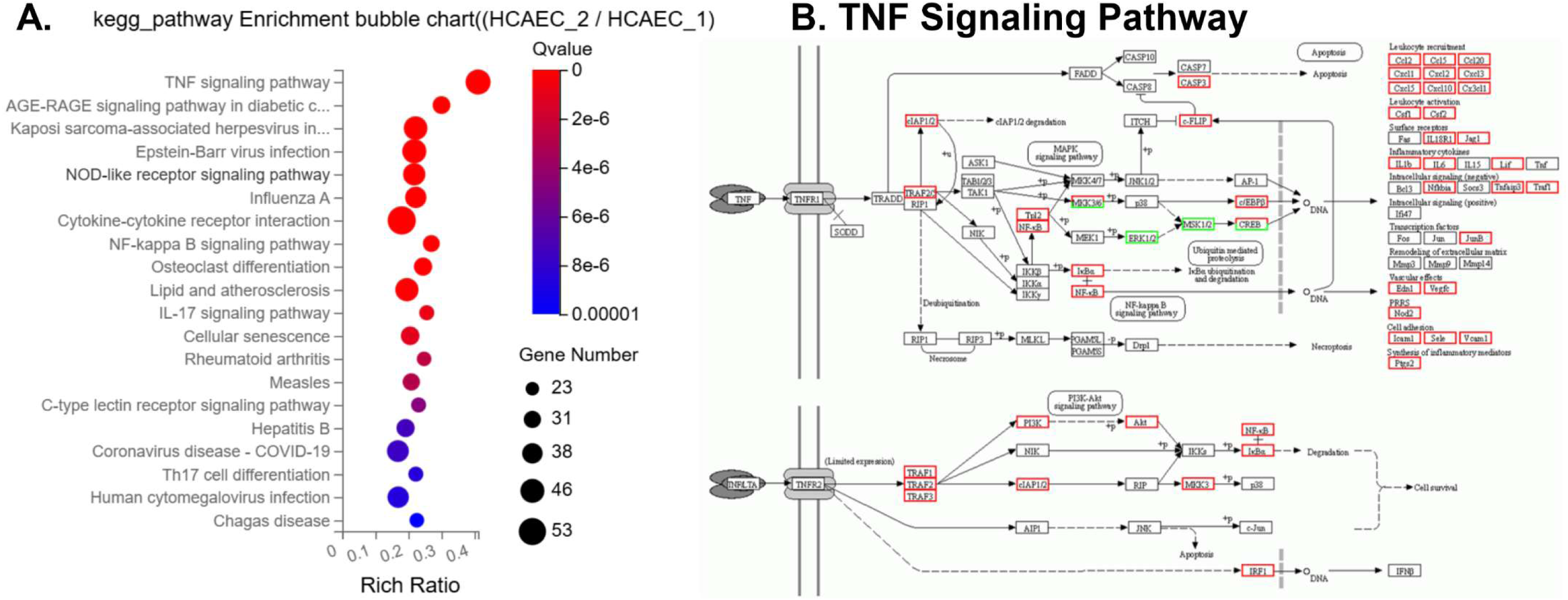
Signaling pathways based on differentially expressed genes induced by medin treatment on HCAECs. A. Kegg pathway analyses show that genes involved in pro-inflammatory signaling were the most affected, with tumor necrosis factor (TNF) and advanced glycation endproducts-receptor for advanced glycation endproducts (AGE-RAGE) signaling as having the highest hits. B shows a preponderance of upregulation over downregulation of genes involved in the TNF signaling pathway. HCAEC1 denotes medin-treated HCAECs and HCAEC2 denotes vehicle-treated HCAECs.

Table 1B shows the top 10 genes downregulated in HCAECs following medin treatment. The top downregulated gene is casein kinase 2 alpha 3, a gene encoding a protein responsible for cell growth, cell cycle, protein phosphorylation and double-strand break repair^37^. Table 1C shows the top genes involved in atherosclerosis and thrombosis differentially expressed following medin treatment. There is upregulation of IL-1β, ICAM-1, MCP-1, PAI-1 and downregulation of thrombomodulin. Supplement Figure 1 shows differentially expressed genes involved in fluid shear stress and atherosclerosis signaling.

### Proinflammatory gene and protein expression

Figure 2A-G shows dose-dependent increases in gene expression of IL-6, IL-8, VCAM-1 and ICAM-1, IL-1β, MCP-1 and TNF-α confirming results of RNAseq analyses. Figure 2H-I shows p-NFκB, but not total NFκB, was increased following 20 hours of medin treatment versus vehicle control. The elevation in p-NFκB with medin treatment was reversed by co-treatment with NFκB inhibitor RO106-9920. Figure 2J-N show medin-induced increases in IL-6, IL-8 and MCP-1 secretion and VCAM-1 and ICAM-1 protein expression were reversed by co-treatment with RO106-9920.

**Figure 2.**
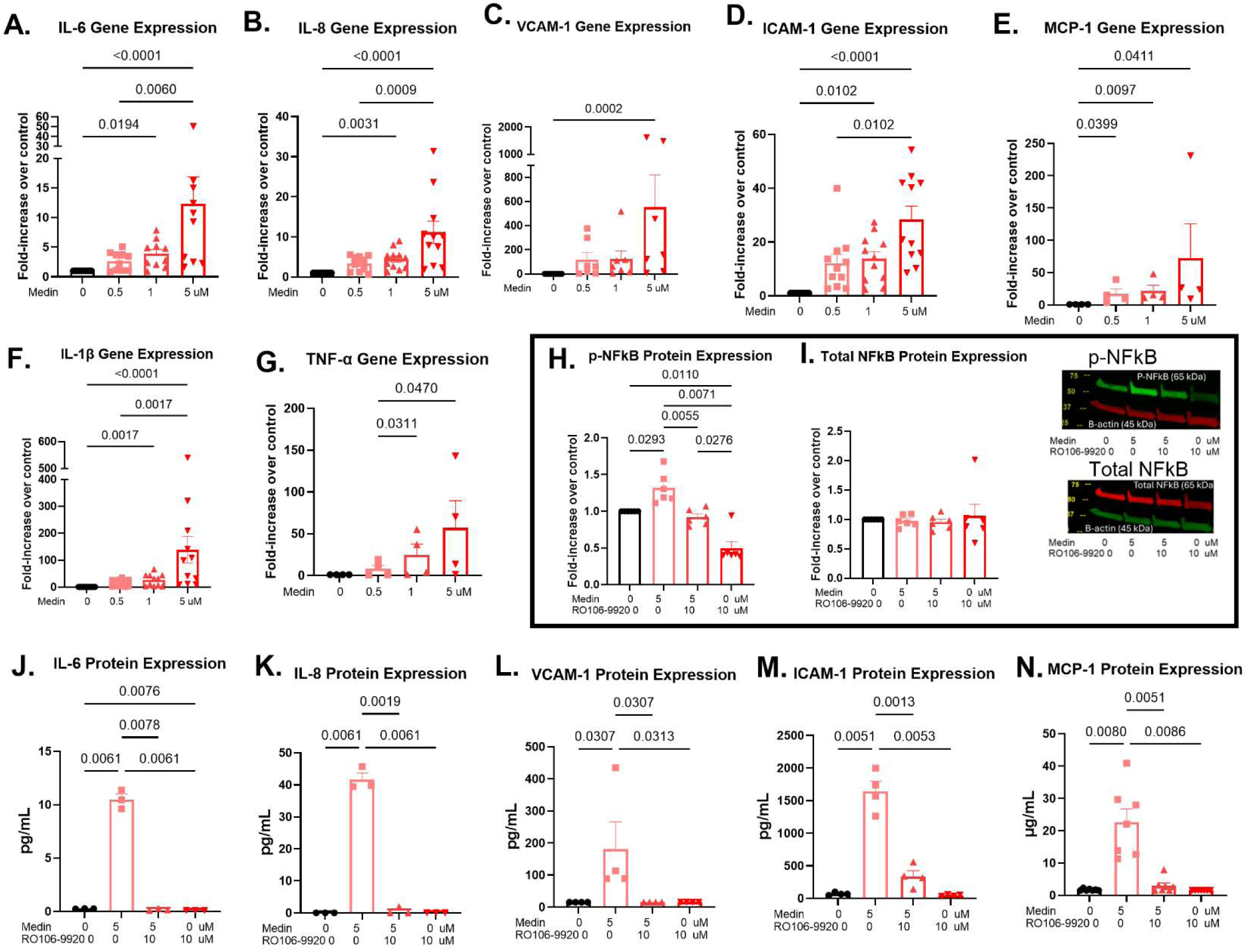
Medin induces pro-inflammatory activation of HCAECs that is NFκB-dependent. A-G show dose-dependent increase in gene expression of IL-6, IL-8, VCAM-1, ICAM-1, IL1β, MCP-1 and TNF-α when HCAECs are treated with 0, 0.5, 1 and 5 µM medin. H shows that after 20 hours of exposure, there remains increased phosphorylated NFκB (p-NFκB) following medin treatment that was reversed by co-treatment of NFκB inhibitor RO106-9920. On the other hand, there was no difference in total NFκB expression (I). Representative Western blots are shown to the right of the graph. J-N show that the protein secretion of IL-6, IL-8 and MCP-1 and the protein expression of VCAM-1 and ICAM-1 in HCAECs are increased by medin 5 µM. These were reversed by co-treatment with RO106-9920. Each datapoint represents a biologic replicate.

To confirm that the effects are due to medin-specific effects rather than non-specific protein effect, separate experiments were done comparing vehicle, medin and scrambled medin. Supplement Figure 2 A-C shows that medin but not scrambled medin increased gene expression of IL-8, ICAM-1, VCAM-1. In similar fashion, medin but not scrambled medin reduced gene expression of thrombomodulin (Supplement Figure 2D). This shows that the effects are specific to medin rather than nonspecific protein effect.

### Prothrombotic gene and protein expression

There was dose-dependent reduction in thrombomodulin gene expression (Figure 3A). Medin also reduced protein expression of thrombomodulin that was reversed by co-treatment of RO16-9920 (Figure 3B). To assess the implication of thrombomodulin reduction in protein C activation, HCAECs exposed to medin were co-treated with thrombin and protein C. Medin reduced activated protein C production that was partly rescued by supplementation with increasing doses of soluble thrombomodulin (Figure 3 C-D) suggesting that medin’s effect on reduced activated protein C is through its downregulation of HCAEC thrombomodulin production.

**Figure 3.**
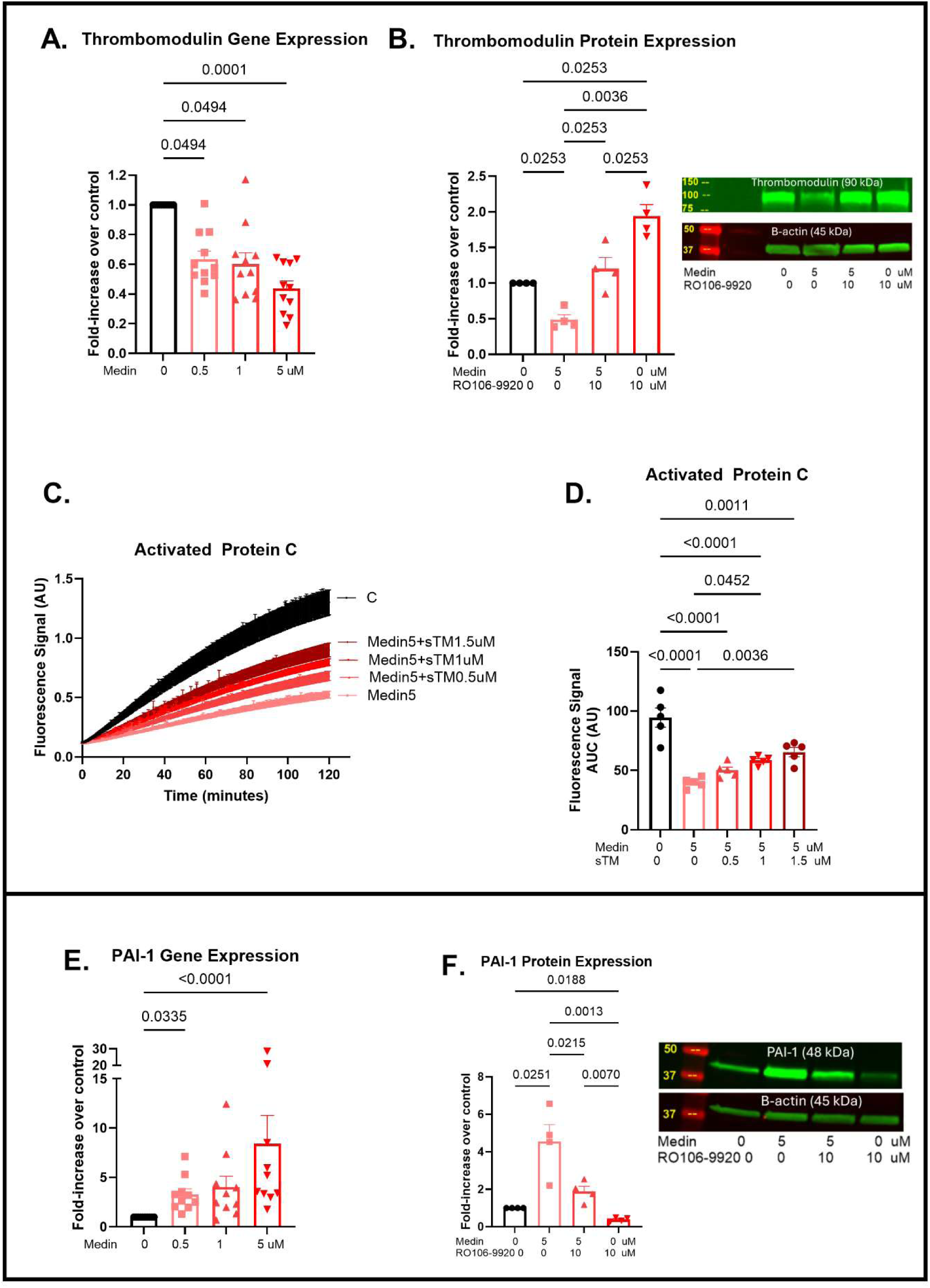
Medin induces prothrombotic activation of HCAECs that is NFκB-dependent. A shows dose-dependent reduction in thrombomodulin gene expression in HCAECs treated with 0, 0.5, 1 and 5 µM medin. B shows that thrombomodulin protein expression was reduced by medin 5 µM but was restored by co-treatment with NFκB inhibitor RO106-9920. C-D shows that HCAECs treated with medin in presence of protein C and thrombin result in reduced activated protein C production when compared with vehicle control but co-treatment with increasing doses of exogenous soluble thrombomodulin restores activated protein C production. This shows that medin’s effect on reducing activated protein C production is due to altered thrombomodulin production. E show dose-dependent increase in PAI-1 gene expression in HCAECs treated with 0, 0.5, 1 and 5 µM medin. F shows that medin-induced increase in HCAEC PAI-1 protein expression is reversed by co-treatment with RO106-9920. Each datapoint represents a biologic replicate.

PAI-1 gene expression had dose-dependent increases following medin treatment (Figure 3E). Medin’s effect on increased PAI-1 protein expression was reversed by co-treatment with RO106-9920 (Figure 3F).

### Human coronary histology

Coronary arteries were analyzed from an original cohort of 14 Arizona decedents (73.6±2.6 years, 9 females, 10 with myocardial infarction history). Figure 4A shows immunohistochemistry with anti-medin monoclonal antibody using DAB staining of coronary arteries that are representative of arterial medin content grouped by quartiles. Figure 4B shows Movat pentachrome staining of the same arteries with clear delineation of the intima/neointima, media and adventitial regions. Note in the examples that medin accumulation is seen especially in complex plaques. Figure 4C-D are magnified images of specific regions. Figure 4E shows CD68 DAB staining of select regions. Overall, coronary medin area was 3.90±1.9% (range 0-39.62%). Plaque burden (area) was 46.96±4.1% (range 14.6-65.5%). CD68+ % area was 0.49±0.26% (range 0-5.6%).

**Figure 4.**
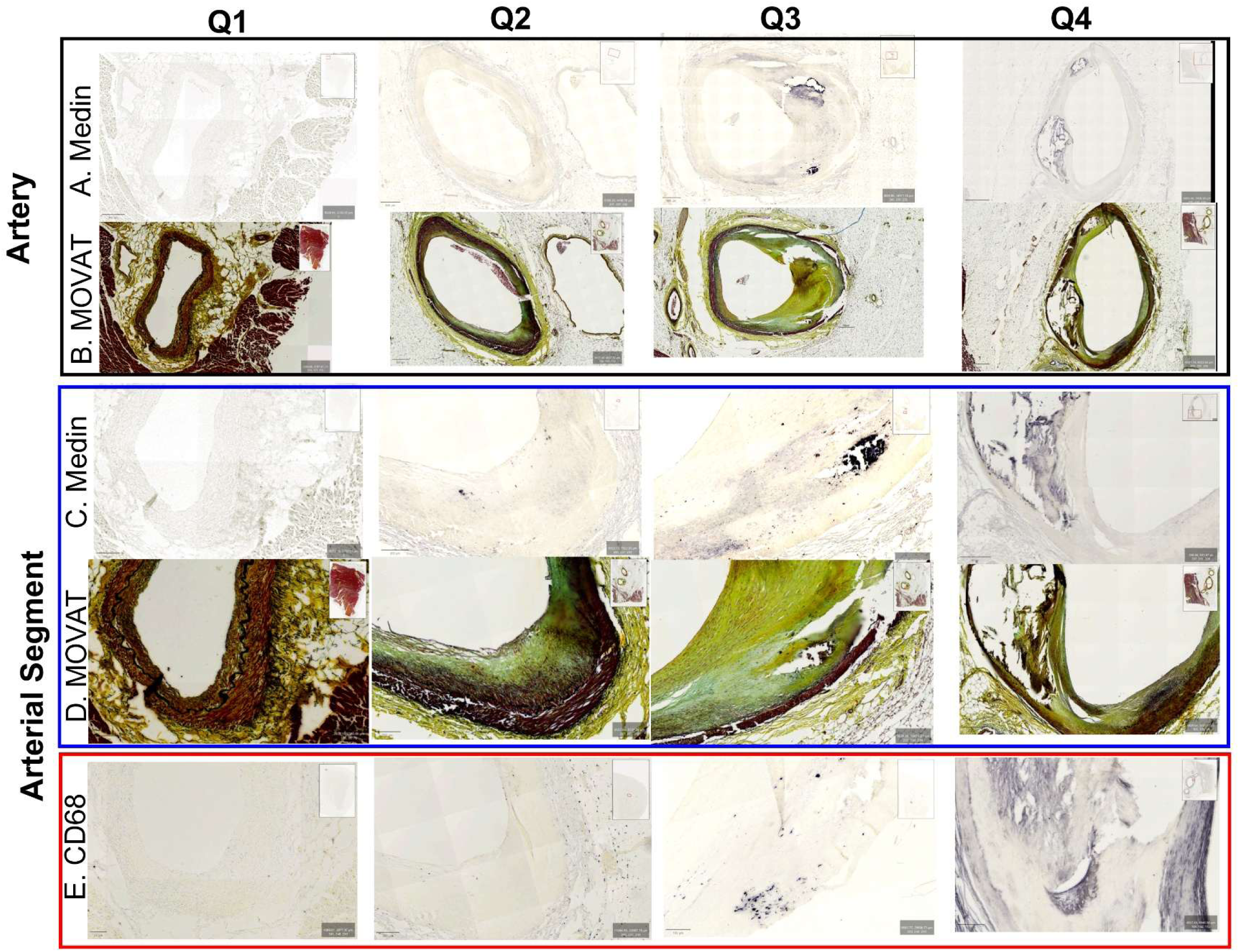
Human coronary medin and plaque. A-B shows epicardial coronary artery sections from donors with DAB immunostaining using anti-medin antibody (A) and Movat pentachrome staining of the same tissue slide (B). C-D are magnified regions of the corresponding arteries in A-B. Each coronary artery represents each quartile (Q1-Q4) by medin area% in the coronary arteries. The images show greater medin (dark blue stain) in coronary arteries with large complex plaques. E shows macrophage DAB immunostaining using anti-CD68 antibody from corresponding regions shown in C-D. Similar to medin staining, there was greater CD68-positive areas with increasing quartile of medin burden.

Medin content as % area of intima-media area was divided into quartiles and the corresponding % plaque area was compared among the quartiles (Figure 5A). There was significantly higher plaque area in the highest quartile of coronary medin compared to the first, second and third quartile groups. In this cohort, there was strong correlation between % medin area and plaque area % (Spearman R=0.72, p<0.0001). Similarly, the macrophage content (CD68+ area %) was higher in the fourth quartile compared to the first and second quartile (Figure 5B); the correlation between the two were significant (Spearman R=0.72, p<0.0001). Coronary medin was significantly higher in coronary arteries from decedents who had myocardial infarction versus those who did not (Figure 5C). A similar although lesser degree of difference was seen for coronary plaque area (Figure 5D).

**Figure 5.**
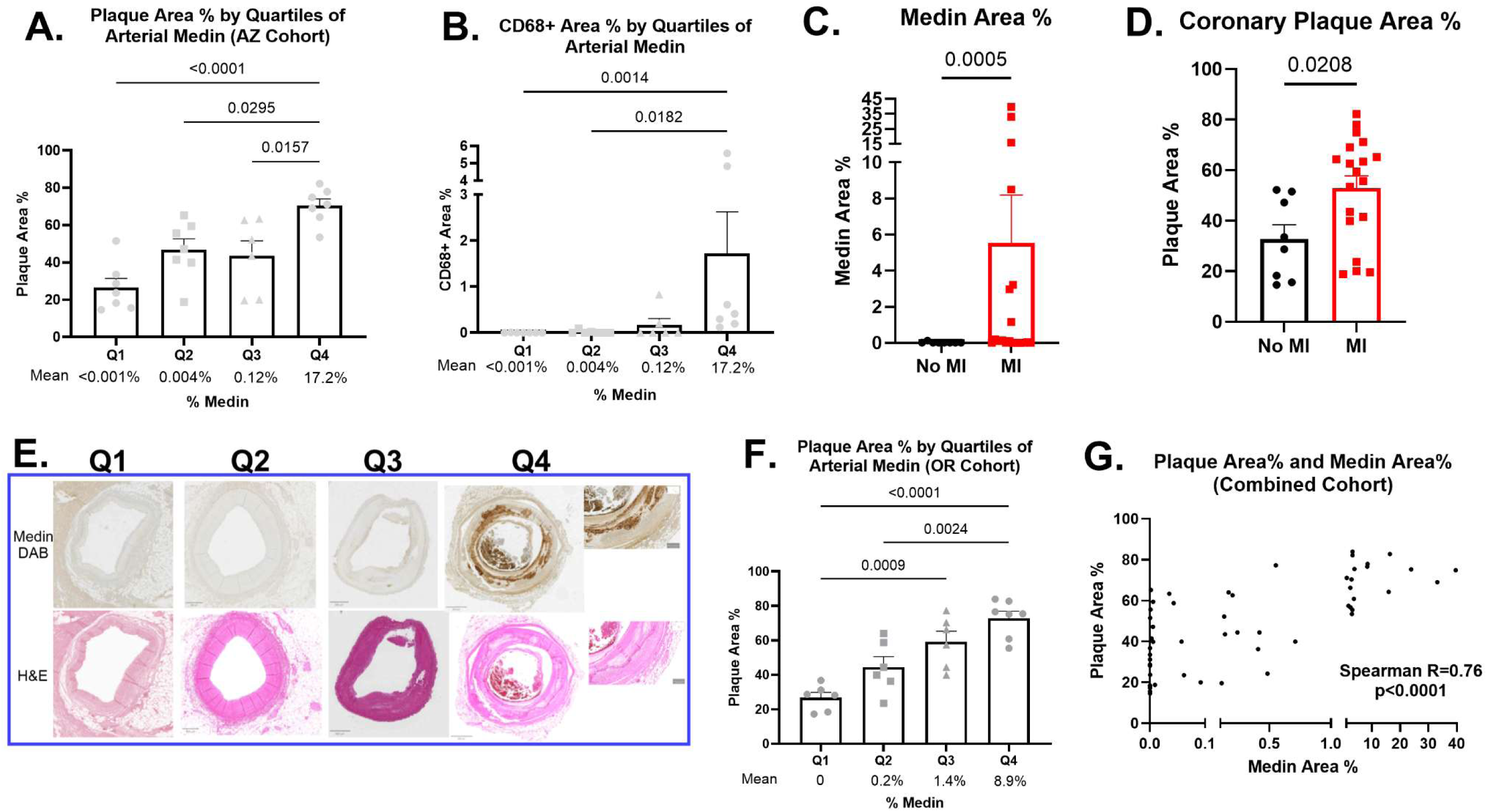
Coronary artery medin burden is related to plaque and macrophage burden and myocardial infarction. A shows the plaque burden (plaque area %) for coronary arteries grouped by quartile of medin burden (medin area %). The highest quartile of medin burden had the highest plaque burden with the lowest quartile having the least amount of plaque burden. B shows the macrophage burden (CD68 area %) grouped by quartile of medin burden. The highest quartile of medin burden had greater macrophage burden compared to the 2 lowest quartiles. C-D show that the coronary medin and plaque burden from donors who had myocardial infarction were higher than donors without myocardial infarction. E-F. A separate cohort of coronary arteries from decedents at OHSU were analyzed, and similar to the Arizona cohort, the plaque burden was higher in coronary arteries with the highest medin burden. G. Combined data from the Arizona and OHSU cohorts show significant correlation between coronary artery medin area% and plaque area%, although the relationship is not linear. Each datapoint represents data from a single coronary artery.

To externally validate our findings, a separate cohort of coronary arteries from 26 decedents at OHSU were analyzed for medin and plaque area (46.6±4.3 years, 11 females). Coronary medin area % for this cohort was 2.89±1.1% (range 0-23.9%) while plaque area % was 51.76±4.2% (range 17.3-83.9%). Similar to the Arizona cohort, there was higher plaque area % with increasing quartile of coronary medin (Figure 5E-F) with significant correlation (Spearman R=0.83, p<0.0001). Similar to the Arizona cohort, there was higher medin in large complex plaques (Figure 5E). When both Arizona and OHSU cohorts were combined, there was strong correlation between coronary medin content and plaque burden (Spearman R=0.76, p<0.0001) (Figure 5G).

## DISCUSSION

The study shows that physiologic doses of medin induce pro-inflammatory and prothrombotic activation of HCAECs that is mediated by NFκB. In human decedents, there was strong correlation between coronary medin content and plaque burden and between coronary medin and macrophage content. There was greater amount of arterial medin in coronary arteries of decedents with MI versus those without. The findings suggest that medin could be a novel mediator of atherosclerosis and could be a critical molecular link between aging and CAD.

Medin was first discovered as the protein composing aortic medial amyloid^11^ and despite being the most common human amyloid which accumulates in the vasculature with advancing age^12,22^, it’s biology remains one of the least known. Medin is a component of the C2 domain of the protein MFGE8 and is cleaved and secreted through still unknown processes and triggers. Arterial medin was found to be higher in cerebral arteries of vascular dementia and Alzheimer’s disease patients when compared to non-demented brain donors^13^ and ex-vivo human cerebral and adipose arteries showed impaired endothelium- and smooth muscle-dependent vasodilation when exposed to physiologic doses of medin as a result of oxidative and nitrative stress^14,15^. A preclinical model of aging C57BL/6 mice showed accumulation of MFGE8 aggregate-like products in the cerebral arteries of 20 month old mice with improved cerebral arterial dilation and constriction in aged mice that had medin (C2 domain) knockout versus controls^20^ validating for the first time the effects of medin on aging-induced vascular dysfunction. In a 2005 study, medin was found to be present in the aorta of all 18 patients age 57 years or older, but in the left coronary artery in only 5 of these patients^12^. The pathologic role of medin in CAD remained unknown and current results provide novel insights.

Medin treatment showed a profound pro-inflammatory activation with increased expression of IL-6, IL-8, IL-1β, VCAM-1, ICAM-1, IL-1β, MCP-1 and TNF-α in HCAECs. Similar results were shown in human brain microvascular endothelial cells^15–17^ and in these prior studies, co-treatment with specific inhibitors showed that medin’s pro-inflammatory activation was mediated by oxidative stress, RAGE and NFκB signaling^15^. Endothelial production of IL-6, IL-8, IL-1β, MCP-1 and TNF-α induce increased vascular permeability and recruitment of inflammatory cells including monocytes/macrophages which in turn results in a feedback loop as the activated macrophages secrete the same cytokines/chemokines, especially TNF-α^38^. Increased adhesion molecules VCAM-1 and ICAM-1 promote leukocyte binding and transmigration^39^. The *in vitro* data combined with our finding of strong correlation between coronary medin content and macrophage burden supports the novel concept of the potential role of medin in coronary arterial inflammation. Chronic low grade vascular inflammation leads to atherosclerotic plaque buildup and our finding that coronary medin content was strongly associated with plaque burden is further support of the possible mediator role of medin in CAD. That the observation of the strong association between medin and coronary plaque burden was seen in both the Arizona and OHSU cohorts support the rigor and robustness of the finding.

In addition to inflammation, aging is associated with the development of a prothrombotic imbalance, and our results suggest that medin could play a role. We previously showed that medin induced increased PAI-1expression in human brain microvascular endothelial cells^16^ and results of the current study shows similar results in HCAECs. PAI-1, a member of the superfamily of serine-protease inhibitors, is the principal inhibitor of both the tissue-type and the urinary-type plasminogen activator. Fibrinolysis results from the interactions among plasminogen activators and inhibitors constituting the enzymatic cascade and ultimately leading to fibrin degradation^40^; increased PAI-1 tends to increase thrombosis and reduce fibrinolysis. PAI-1 expression, like medin production, is increased in the elderly^41^. Our results suggest that medin could be one of the mediators of increased PAI-1 with aging.

Another novel finding of our study shows that medin induced reduction in HCAEC gene and protein expression of thrombomodulin. Thrombomodulin is a thrombin receptor on endothelial cells that alters the active site specificity of thrombin facilitating the proteolytic activation of protein C which in turn degrades factors Va and VIIIa, inhibiting the clotting mechanism and suppressing further thrombin generation^9,42^. The binding of thrombin to thrombomodulin not only results in an ∼1000-fold increase in activation of protein C but also blocks the thrombin-mediated conversion of fibrinogen into fibrin and inhibits thrombin binding to receptors on platelets and inflammatory cells^43^. The adverse biologic consequence of the reduced thrombomodulin expression induced by medin was confirmed by showing that in the presence of thrombin and protein C, there was reduced activated protein C and that this reduction is a result of reduced HCAEC thrombomodulin since introduction of exogenous soluble thrombomodulin restored activated protein C levels.

The combined reduction of thrombomodulin and the increase in PAI-1 expression in HCAECs treated with medin therefore constitute a prothrombotic transformation that promotes the clotting mechanism (reduced restraint on thrombin, factors Va and VIIIa from reduced thrombomodulin) and reduces fibrinolysis (through increased PAI-1). These *in vitro* data are consistent with our finding that coronary arteries from decedents with MI had greater medin compared to those from decedents without MI. Myocardial infarction is most commonly caused by thrombotic occlusion of a disrupted coronary plaque impairing downstream perfusion. These findings offer support for the possible mediator role of medin in coronary thrombosis.

We previously showed that the pro-inflammatory activation induced by medin on human brain microvascular and human umbilical vein endothelial cells was completely abolished by co-treatment of small molecule specific NFκB inhibitor RO106-9920^15,16^. Our results show similar results in HCAECs. RO106-9920 is a potent inhibitor of NFκB by selectively blocking ubiquitination activity associated with lipopolysaccharide and TNF-α induced IκBα degradation and NFκB activation^44^. Interestingly, medin-induced altered PAI-1 and thrombomodulin expression in HCAECs were also reversed by NFκB inhibition, showing common signaling mechanism between pro-inflammatory and prothrombotic processes with NFκB as the master regulator. NFκB signaling has a key role in endothelial cells in response to stress situations as it is capable of regulating both pro-inflammatory and prothrombotic responses which are also prone to a significant level of bidirectional cross-talk^43^. Specifically, in mice subjected to endotoxemia, endothelial NFκB activation impaired thrombomodulin-protein C anticoagulation pathway^45^. IL-1β expressed following NFκB activation result in increased PAI-1^46^. NFκB-mediated reduction in thrombomodulin leads to reduced activated protein C which is involved in cytoprotection through protease-activated receptor1 (PAR1) signaling that inhibits NFκB activation^42^, linking the clotting and inflammatory cascade in an amplifying feed-forward loop. On the other hand, a negative feedback loop could also play a role, as PAI-1 was shown to inhibit endotoxin-induced TNF-α production by monocytes^43^. The dual role of medin in pro-inflammatory and prothrombotic activation of HCAECs through NFκB signaling could represent a treatment target and novel mediator/risk factor for CAD and myocardial infarction.

The study has several limitations. The study focused on the effects of medin on HCAECs but recent study showed that medin also induced pro-inflammatory activation of human brain vascular smooth muscle cells without affecting phenotypic transformation^47^. Future studies should investigate the effects of medin on coronary artery smooth muscle cells and their role in atherosclerosis. Our data from human tissues show an association between medin and plaque burden, macrophage burden and myocardial infarction but causal relationship cannot be established and only suggested by the supporting plausible mechanistic *in vitro* data presented. Follow-up preclinical studies with knockout or overexpression of medin to see the effects on coronary atherosclerosis should be performed. Finally, the study focused only on select pro-inflammatory and prothrombotic mediators but the plethora of data from the RNAseq analyses should encourage further exploration of other identified altered pathways.

## ACKNOWLEDGMENTS

Funding was provided by VA Merit 1I01BX006216, BX003767 and NIH 1R56AG083570, NIARO1 AG019795, NIAP30AG19610, NINDSU24NS072026) and Michael J. Fox Foundation. We thank Dr. Thomas Beach, Dr. Geidy Serrano and the Banner Sun Health Research Institute Brain and Body Donation Program. We thank Ms. Gail Farrell for regulatory support and the Phoenix VA Office of Research and Arizona Veterans Research and Education Foundation for their logistical support.

The content and views of this manuscript do not reflect the views of the US Department of Veterans Affairs or the United States government.

## Supplement Figures Legend

**Supplement Figure 1.**
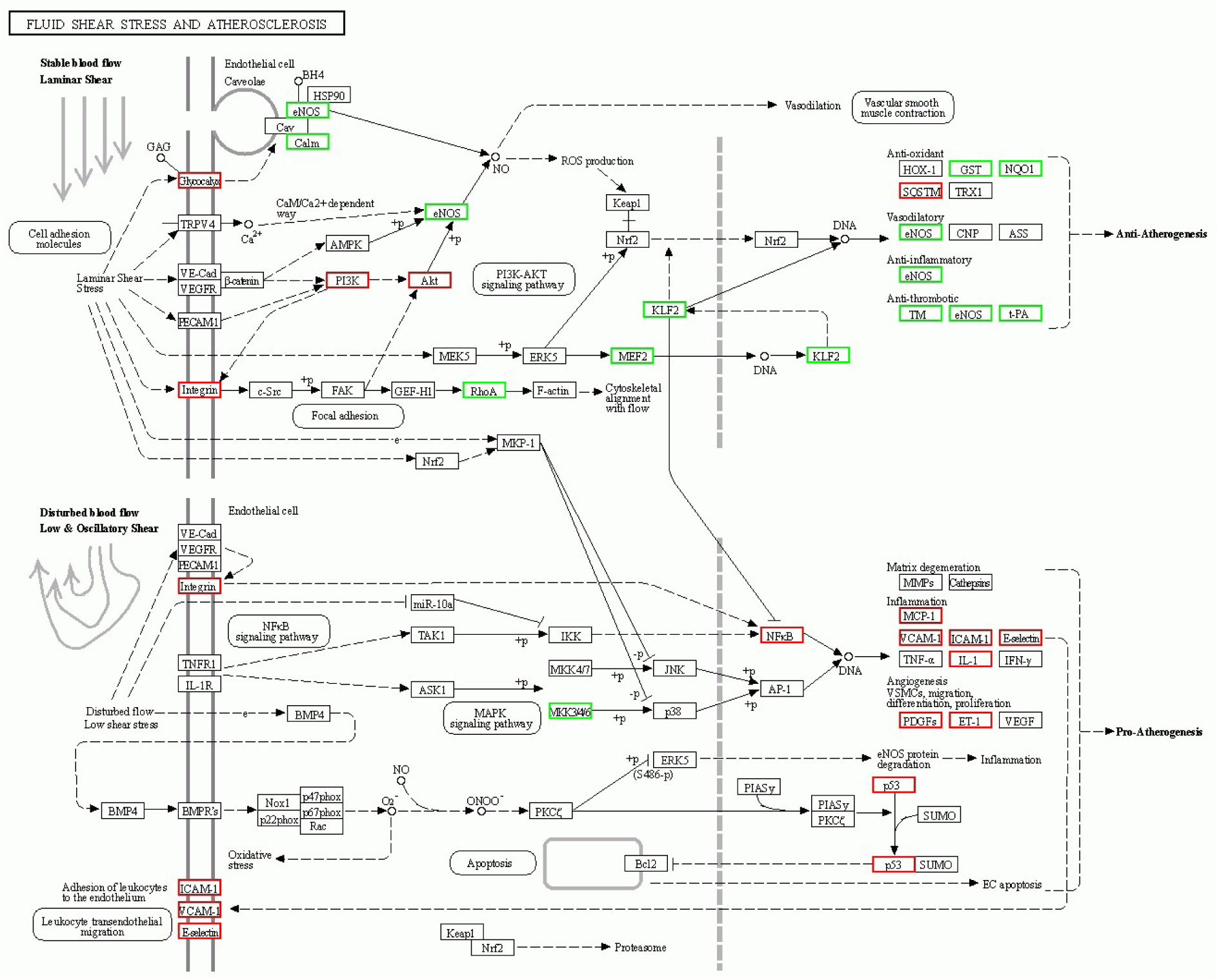
RNA sequencing genes affected by medin treatment of HCAECs associated with the fluid shear stress and atherosclerosis signaling pathway. Downregulated genes are in green and include thrombomodulin and upregulated genes are in red such as VCAM-1 and ICAM-1.

**Supplement Figure 2.**
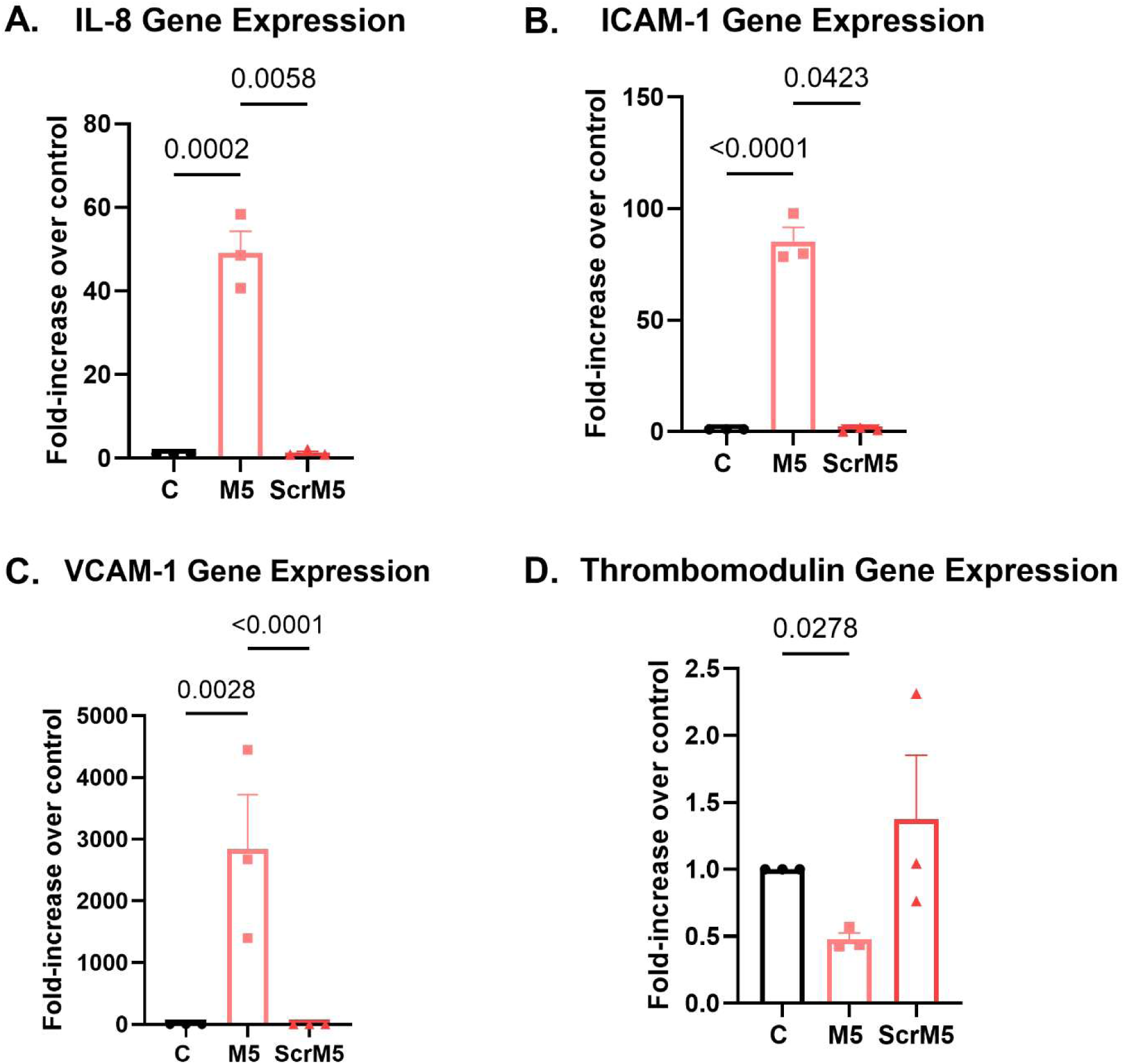
Medin, but not scrambled medin, induces HCAEC activation. A-D shows that HCAECs treated with medin showed increased gene expression of IL-8, ICAM-1 and VCAM-1 and reduced gene expression of thrombomodulin when compared to vehicle control. These effects were not seen with exposure to scrambled medin protein. Each datapoint is a biologic replicate.

